# Selective Maintenance of Value Information Helps Resolve the Exploration/Exploitation Dilemma

**DOI:** 10.1101/195453

**Authors:** Michael N. Hallquist, Alexandre Y. Dombrovski

**Affiliations:** Penn State University, Department of Psychology; University of Pittsburgh, Department of Psychiatry

## Abstract

Laboratory studies of value-based decision-making often involve choosing among a few discrete actions. Yet in natural environments, we encounter a multitude of options whose values may be unknown or poorly estimated. Given that our cognitive capacity is bounded, in complex environments, it becomes hard to solve the challenge of whether to exploit an action with known value or search for even better alternatives. In reinforcement learning, the intractable exploration/exploitation tradeoff is typically handled by controlling the temperature parameter of the softmax stochastic exploration policy or by encouraging the selection of uncertain options.

We describe how selectively maintaining high-value actions in a manner that reduces their information content helps to resolve the exploration/exploitation dilemma during a reinforcement-based timing task. By definition of the softmax policy, the information content (i.e., Shannon’s entropy) of the value representation controls the shift from exploration to exploitation. When subjective values for different response times are similar, the entropy is high, inducing exploration. Under selective maintenance, entropy declines as the agent preferentially maps the most valuable parts of the environment and forgets the rest, facilitating exploitation. We demonstrate *in silico* that this memory-constrained algorithm performs as well as cognitively demanding uncertainty-driven exploration, even though the latter yields a more accurate representation of the contingency.

We found that human behavior was best characterized by a selective maintenance model. Information dynamics consistent with selective maintenance were most pronounced in better-performing subjects, in those with higher non-verbal intelligence, and in learnable vs. unlearnable contingencies. Entropy of value traces shaped human exploration behavior (response time swings), whereas uncertainty-driven exploration was not supported by Bayesian model comparison. In summary, when the action space is large, strategic maintenance of value information reduces cognitive load and facilitates the resolution of the exploration/exploitation dilemma.

**Author summary:** A much-debated question is whether humans explore new options at random or selectively explore unfamiliar options. We show that uncertainty-driven exploration recovers a more accurate picture of simulated environments, but typically does not lead to greater success in foraging. The alternative approach of mapping the most valuable parts of the world accurately while having only approximate knowledge of the rest is just as successful, requires less representational capacity, and provides a better explanation of human behavior. Furthermore, when searching among a multitude of response times, people cannot indefinitely maintain information about every experience. A good strategy for someone with limited memory capacity is to selectively maintain a valuable subset of options and gradually forget the rest. In simulated worlds, a player with this strategy was as successful as a player that represented all previous experiences. When learning a time-varying contingency, humans behaved in a manner consistent with a selective maintenance account. The amount of information retained under this strategy is high early in learning, encouraging exploration, and declines after one has discovered valuable response times.

## Introduction

> *It is better to understand a little than to misunderstand a lot.*
>
> - Anatole France

Laboratory studies of value-based decision-making typically involve choosing among a few actions according to their perceived subjective value (1). In real life, however, we often face a multitude of options whose values may be unknown or poorly estimated. How can an organism with limited computational resources learn the most advantageous actions in the natural environment? Previous work on boundedly rational agents has considered the role of a limited-capacity working memory system (2) and the possibility that metareasoning (i.e., a policy guiding *how* to allocate resources) reduces the complexity of learning in large action spaces (3). This study provides a new, complementary account highlighting how the selective maintenance of value information facilitates the search for the best among many actions.

One of the fundamental dilemmas in reinforcement learning is how to choose between exploiting an action with a known positive value and exploring alternatives in search of even more advantageous actions (4). A much-debated question is whether human exploration is driven by uncertainty (5). An influential idea from artificial intelligence is that agents may receive ‘exploration bonuses’ for exploring highly uncertain states (6), yet studies using multi-armed bandit tasks have not found evidence of this (7). Rather, humans appear to become averse to uncertainty as the number of options increases (8) unless uncertainty-driven exploration is explicitly encouraged (9). On the other hand, Frank and colleagues presented evidence of spontaneous uncertainty-driven exploration on an instrumental reinforcement-based timing task — the clock task — using their Time-Clock (TC) computational model (10,11). Thus, an important unanswered question is whether the large action space of a timing task elicits uncertainty-based exploration even if simpler discrete choice paradigms do not.

In reinforcement-based timing tasks, optimal timing is often uncertain, but can be learned by responding at different moments in time and evaluating the outcome. Unlike learning tasks with just a few actions, reinforcement-based timing requires one to explore a continuous action space to identify response times with high expected value. Assuming some degree of temporal generalization, the complexity of representing time-varying reinforcement can be reduced by a set of temporal basis functions (TBF) that approximate expected value as a function of time. A TBF representation has been validated in temporal difference (TD) models of Pavlovian conditioning (12,13), providing a parsimonious account of timing that limits the number of values maintained in memory. A further challenge, however, is that memory traces inevitably decay over time, particularly when many action values are kept online (14,15). Thus, effective approaches to learning need to be robust to forgetting, ensuring that valuable information is selectively maintained (cf. 16). Building on models of working memory (17,18) and the dopamine system (19), we propose that in reinforcement-based timing, the values of recently sampled actions (response times) are selectively maintained, whereas more temporally distant action values decay. As we illustrate below, such selective maintenance trades off a high-fidelity representation of all available rewards for the opportunity to exploit the best response time (cf. 20).

Algorithmic solutions to the intractable exploration/exploitation dilemma generally focus on the policy, which translates subjective value estimates into choices. The commonly used Boltzmann softmax policy probabilistically selects actions in proportion to their value, while controlling exploration with a temperature parameter (4). The degree of exploration in a given state depends on the *entropy* of the Boltzmann distribution of action probabilities, a logarithmic measure of uncertainty about which action to choose (21). Entropy is maximal when all actions have equal probability and approaches zero as the probability of choosing one action approaches one. In turn, when the temperature parameter is high, entropy increases and actions are chosen with similar probabilities (exploration), whereas if it is low, the agent prefers high-value actions (exploitation).

While prior work has focused on controlling the softmax temperature or encouraging uncertainty-driven exploration, the similarity among the learned action values *per se* is a more fundamental determinant of the exploration/exploitation tradeoff. We quantify this similarity as Shannon’s entropy (22) of the normalized vector of action values (cf. 8), which represents the information content of the learned values. For example, if the expected value of one action is much greater than the alternatives, the entropy of action values will be low, increasing the likelihood of exploiting the valued action in the softmax policy. Below, we show that in the case of reinforcement-based timing, selective maintenance of action values across learning episodes helps to resolve the exploration/exploitation dilemma by reducing entropy. We demonstrate *in silico* and *in vivo* that selectively maintaining values of recently chosen actions while gradually forgetting temporally remote ones is generally superior to tracking the value of all available actions.

Altogether, these considerations motivated the hypothesis that humans selectively maintain value traces in a manner that compresses them, reducing memory load. Second, we hypothesized that the resulting entropy dynamics tune the exploration/exploitation tradeoff via the Boltzmann softmax, better accounting for human sampling behavior than directed exploration toward uncertain options (6,11,23). To test these hypotheses, we developed a new reinforcement learning model, StrategiC ExPloration/ExPloitation of Temporal Instrumental Contingencies (SCEPTIC), that represents continuous action values using basis functions. We tested variants embodying alternative hypotheses, using the temporal difference (TD) model as a benchmark and Frank’s TC as a comparator. In model comparisons, we used model and parameter identifiability and optimality as preliminary criteria and fit to behavior as the final criterion. We took particular care to rule out alternative accounts of behavior and information maintenance, such as exploration driven by local uncertainty, variable learning rate, and choice autocorrelation.

## Results

### The clock paradigm and model-free overview of behavior

The clock task is depicted in Figure 1a-b. Subjects were asked to find the “best” response time during a 4 s interval. Outcomes were controlled by one of the four probabilistic contingencies with varying probability/magnitude tradeoffs (1b), two of them learnable with value maxima in the beginning (decreasing expected value, DEV) or the end (increasing expected value, IEV) of the interval, and two unlearnable (constant expected value, CEV and constant expected value-reversed, CEVR). This design results in a high level of uncertainty and encourages trial-by-trial learning, making it difficult to find an optimal strategy.

**Figure 1.**
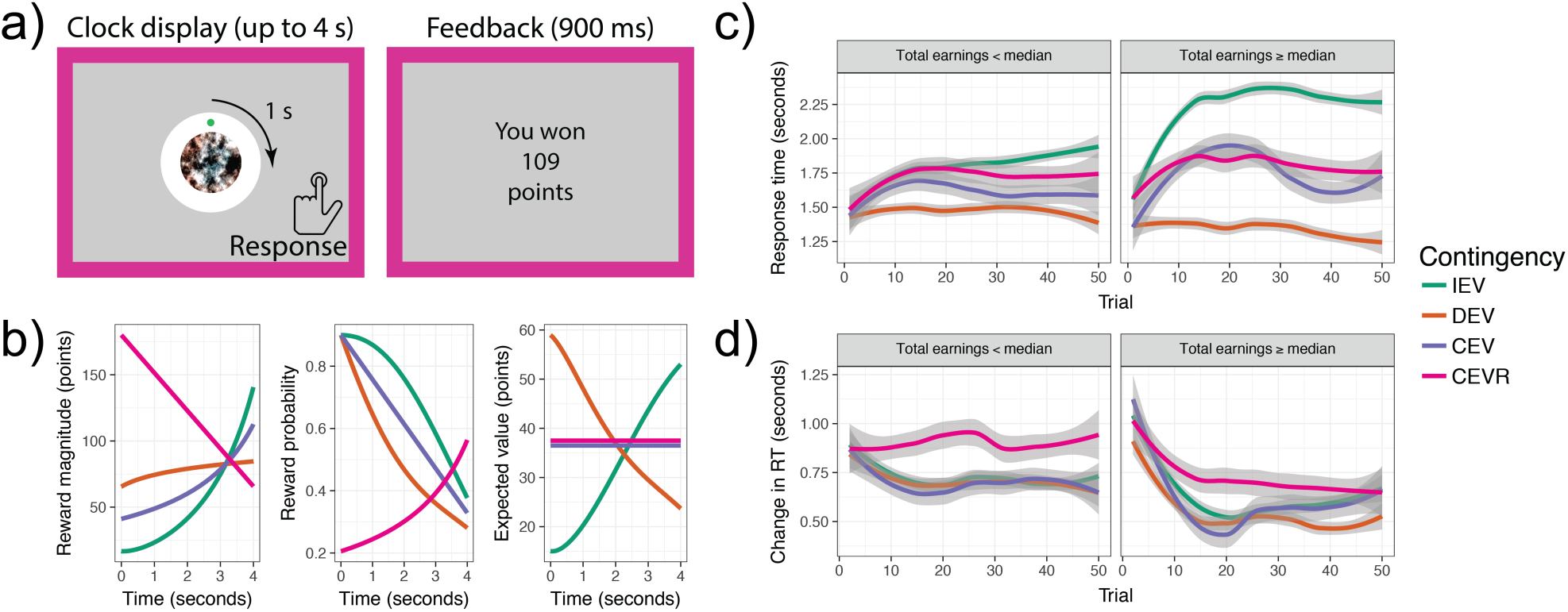
The clock paradigm and typical human behavior. a) The clock paradigm consists of decision and feedback phases. During the decision phase, a dot revolves 360° around a central stimulus over the course of four seconds. Participants press a button to stop the revolution and receive a probabilistic outcome. b) Rewards are drawn from one of four monotonically time-varying contingencies: increasing expected value (IEV), decreasing expected value (DEV), constant expected value (CEV), or constant expected value–reversed (CEVR). CEV and CEVR thus represent unlearnable contingencies with no true value maximum. Reward probabilities and magnitudes vary independently. c) Evolution of subjects’ response times (RT) by contingency and performance. Panels represent participants whose total earnings were above or below the sample median. d) Evolution of subjects’ response time swings (RT swings) by contingency and performance.

As in previous studies (10,24), with learning, subjects’ response times (RT) rapidly shifted toward value maxima: late in the interval in IEV and very early for DEV (Figure 1c). As expected, these shifts were more prominent in better-performing subjects and were not apparent in unlearnable contingencies. The rate of exploration, as measured by trial-wise change in response times (i.e., ‘RT swings’), declined with learning and was higher in unlearnable contingencies (CEV, CEVR; Figure 1d) and also in poorly performing subjects (Fig. 1d, right vs. left panel), highlighting a stochastic underlying process. Interestingly, RT swings declined in both learnable and unlearnable contingencies. Even more remarkable was the fact that the switch from exploration to exploitation in both unlearnable contingencies was more pronounced in better-performing subjects, indicating that they tended to settle into a perceived value maximum even where objectively there was none. This suggests that successful learners rely on a mechanism that accelerates the transition from exploration to exploitation. At the same time, these results cast doubt on the strategic uncertainty-driven nature of RT swings, at least beyond the first few trials, for two reasons. First, uncertainty-driven exploration should improve performance by uncovering a value maximum, while we see the exact opposite: persistent RT swings reflect stochastic responding and indicate ignorance of the value maximum in learnable contingencies. Second, uncertainty-driven exploration cannot explain higher RT swings in unlearnable contingencies,since constant expected value leads to more uniform sampling (Figure 1c), which diminishes uncertainty gradients. In summary, stochastic exploration appears to underlie RT swings without an obvious need to invoke uncertainty-seeking.

### SCEPTIC architecture (Table 1)

Unlike tasks with a few actions whose values can be learned discretely, the clock task has a rather large action space in which the expected value of responding at any particular moment may be unique (Figure 1a-b). The SCEPTIC model approaches this as a function approximation problem in which an agent with limited cognitive resources and an imprecise representation of time learns a heuristic representation of expected value (cf. 25). Extending earlier work on time representation in Pavlovian learning (12), SCEPTIC uses Gaussian TBFs to approximate the time-varying contingency. Each function has a temporal receptive field with a mean and variance defining its point of maximal sensitivity and the range of times to which it is sensitive. The weights of each TBF are updated according to a delta learning rule (Figure 2; detailed in Materials and Methods). Further, whereas learning and choice in TD occur on a moment-to-moment basis, humans are often more strategic, considering the entire interval at once, an observation reflected in Frank’s TC model (11). Building on this insight, SCEPTIC considers updates and choices at interval level. All SCEPTIC models (Table 1) shared this general architecture.

**Table 1:** Summary of key models tested

| Model |  | Temporal basis function | Uncertainty-driven exploration | Value learning rule |  | Interval-level (vs. time step-level) policy | Response | Number of free parameters |
| --- | --- | --- | --- | --- | --- | --- | --- | --- |
|  |  |  |  | Delta rule with fixed learning rate | Kalman filter |  |  |  |
| SCEP TIC | Fixed learning rate, value (LR V) | + | - | + | - | + | Multinomial | 2 |
|  | Fixed LR uncertainty + value (U+V) | + | + | + | - | + |  | 3 |
|  | Fixed LR selective maintenance | + | - | + | - | + |  | 3 |
|  | Kalman filter (KF), softmax | + | - | - | + | + |  | 1 |
|  | KF U+V | + | + | - | + | + |  | 2 |
| Time-clock (TC) |  | - | + | + | - | + | Continuous | 7 |
| Temporal difference (TD), Q-learning |  | - | - | - | - | - | Multinomial | 2 |

**Figure 2.**
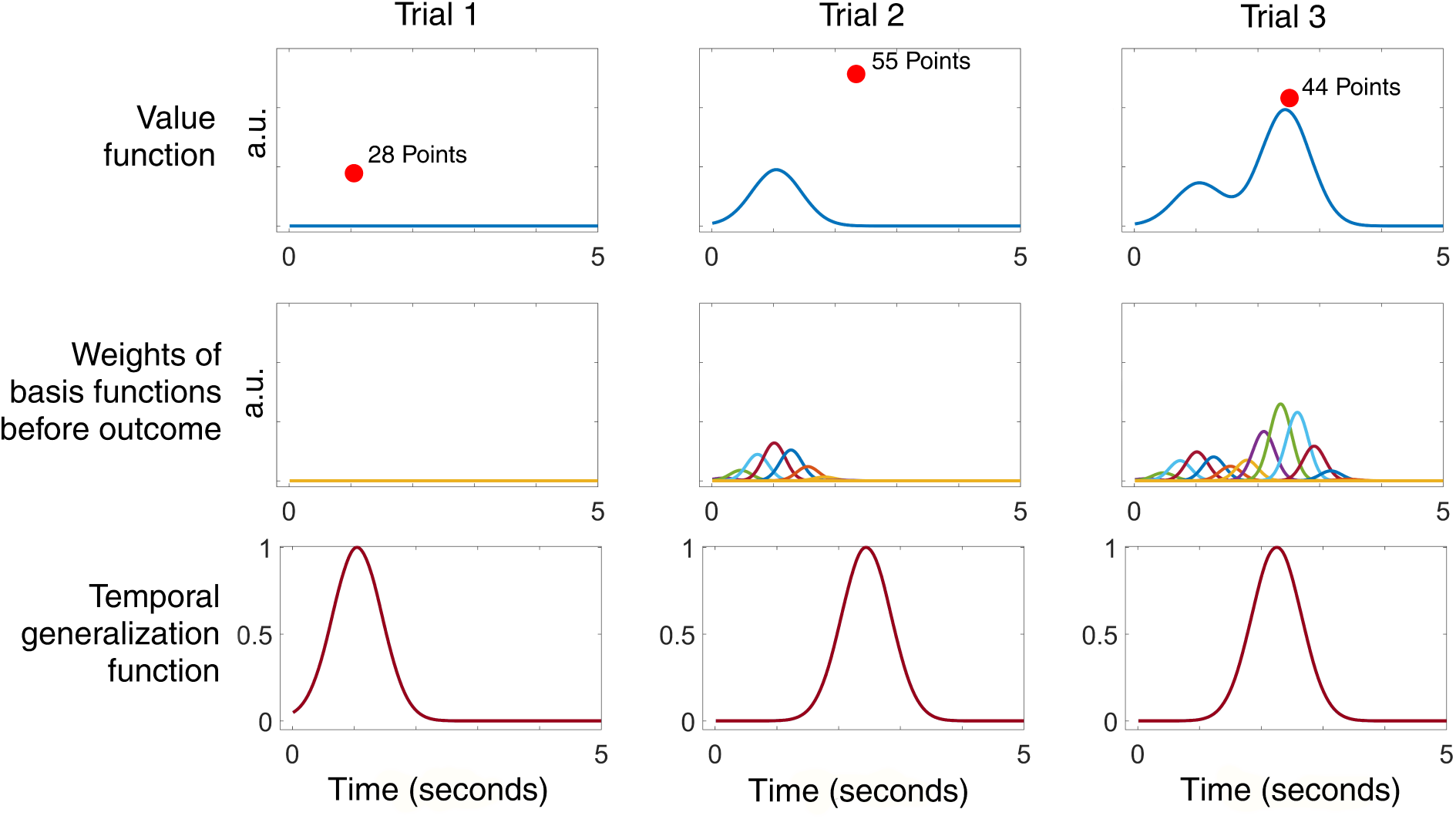
The SCEPTIC model represents the clock paradigm using a set of temporal basis functions (TBFs) spaced evenly over the time interval (middle row). These TBFs approximate a continuous time-varying expected value function (top row). The effect of prediction errors on value estimates are spread symmetrically in time according to a temporal generalization function centered on the chosen response time (bottom row). The red dot represents the chosen response time and the adjacent text indicates the reward outcome. Data from a randomly selected subject. AU = arbitrary units.

#### SCEPTIC: selective maintenance

To test the hypothesis of selective maintenance of value representations under cognitive constraints, we developed a version of the SCEPTIC model, Fixed LR V selective maintenance, that allowed for the forgetting of value traces that were not selected on a given trial (illustrated in Figure 3). Its reward values reverted toward zero in inverse proportion to a temporal generalization function (Figure 2). Below, we refer to it as the “selective maintenance model”. By erasing the reinforcement history in seldom-visited parts of the interval, such selective maintenance tends to decrease the information content of the action value representation later in learning (Fig 3b). We quantified the amount of information contained in the value representation as Shannon’s entropy of the vector of basis function weights, **w**. The weights were first normalized to have an area under the curve of unity (cf. 26), although we note that other normalization methods will yield similar results.

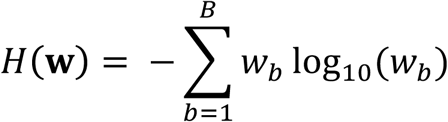

**Figure 3.**
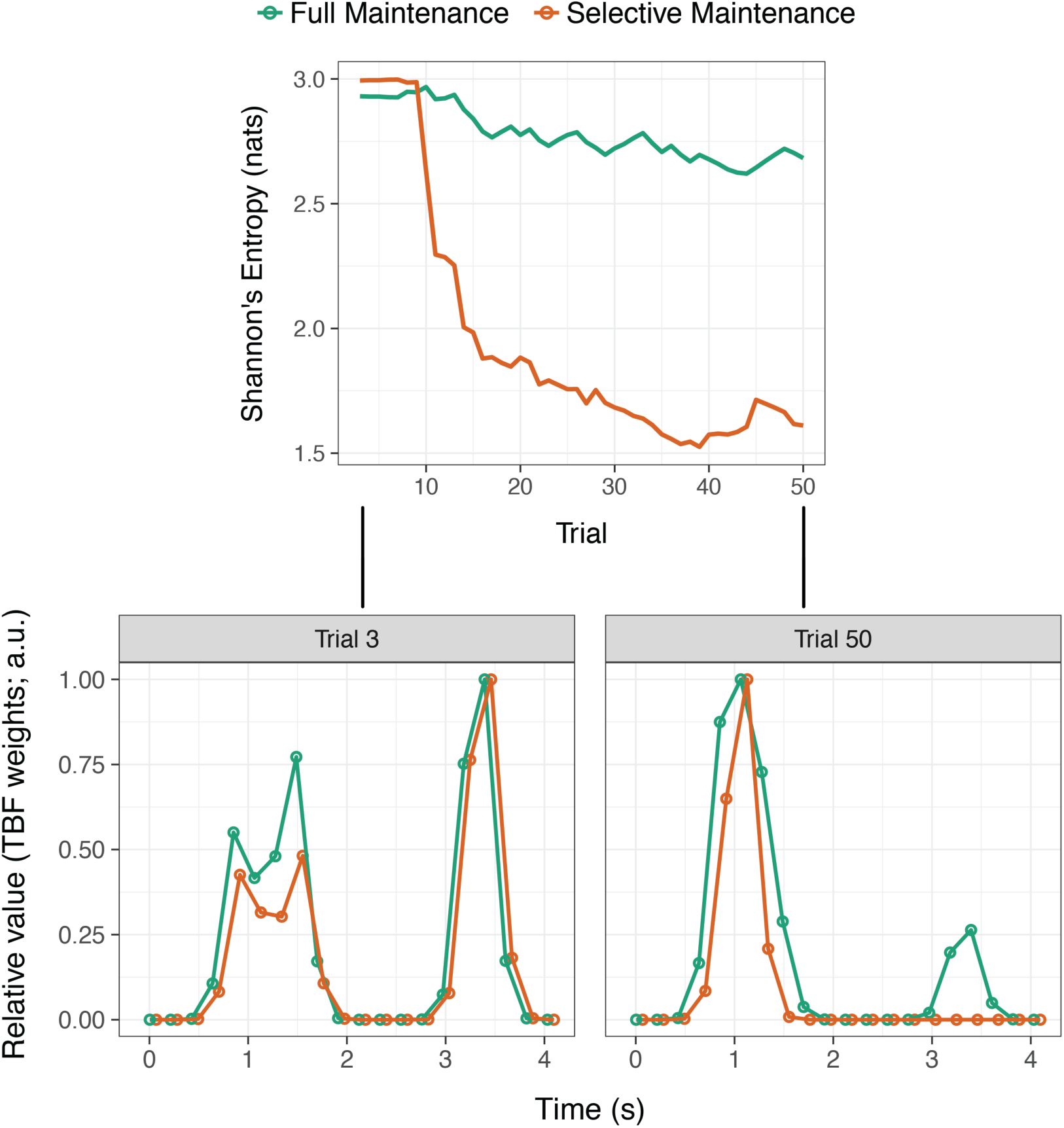
Representation of expected value in SCEPTIC models with and without selective maintenance (data from a representative subject). Top: evolution of entropy over trials; bottom left: action values at trial 3; bottom right: action values at trial 50. Notable differences emerge late in learning (right). Whereas the full-maintenance model (Fixed LR V; green line) contains a more detailed representation of the contingency, the selective maintenance model (orange line) tends to track a single value bump corresponding to a hypothesis about the best response time. This corresponds to a lower information content (entropy) of value representations in the selective maintenance model compared to the full model (see Figure 7 for more detail). The dots in the figure denote the weights for each basis function, which are multiplied by the Gaussian basis to form the integrated value representation, *V* (*i*). Although the agent selects actions based on *V*(*i*), we depict basis weights here because they are used to compute entropy, as shown in the top panel. The maximum of each representation is rescaled to the same value (1.0) to facilitate comparison. Absolute values will be depressed by selective maintenance, but only relative values impact choice. a.u. = arbitrary units.

### SCEPTIC: the impact of uncertainty on exploration

To embody the alternative hypothesis that exploration is modulated by uncertainty, we developed SCEPTIC variants where choice was influenced by both *uncertainty* (U), estimated by Bayesian filtering, and reward *value* (V). In U+V models, choice was controlled by a weighted sum of uncertainty and value according to a parameter, *τ* uncertainty and value according to a parameter, that could assume positive values reflecting uncertainty-driven exploration, or negative values reflecting uncertainty aversion. Since uncertainty may impact not only exploration but also the learning rate (25–28), we examined the impact of uncertainty in both fixed learning rate (LR) models and in models where the learning rate was controlled by a Kalman filter (KF). Further, to ascertain that our model comparison results are not limited to this specific implementation of uncertainty-driven exploration and fixed or dynamic learning rate, we tested a number of alternative models, described in the Supplementary Materials.

### Alternative account of uncertainty-driven exploration: the time-clock (TC) model

Developed by Frank and colleagues for the clock task (11), TC represents response times as a function of seven decision signals, including three free parameters of no interest reflecting subject’s mean response time, choice autocorrelation (27), and modulation toward the best outcome experienced thus far. Two parameters represent speeding or slowing of response times due to prediction errors, inspired by a computational model of the basal ganglia (28). Two final parameters represent the influence of expected value and outcome uncertainty. One noteworthy aspect of the TC model is that instead of separating the learning rule from the choice rule, decision signals contribute additively to the predicted response time.

### Benchmark: modified temporal difference (TD) model

TD has previously been shown to fail on the clock task (24) due to an erroneous back-propagation of value. To obtain a robust benchmark, we adapted a TD *Q*-learning model with a complete serial compound stimulus representation. As detailed in Materials and Methods, two modifications were needed to improve the performance of TD. First, to encourage the agent to sample the entire time interval, we modified the γ-greedy choice rule to make responding less likely for early than late time steps. Second, to overcome the shift of estimated value from later to earlier regions of the interval, the value update for a reinforced response was back-propagated only to preceding *wait* actions, crediting them appropriately for the reward. Lastly, we compared *Q*-learning and SARSA variants of TD in simulations and participant behavior, finding *Q*-learning to be superior (data available upon request).

### Validation of models in simulated environments

In order to examine whether model parameters could be estimated reliably from behavioral data, we conducted a series of parameter recovery simulations. In a variety of environments, the parameters of key SCEPTIC models (Table 1) were identifiable (all *R* ^2^> .93), whereas the TD and TC models had problems with parameter indeterminacy (see Supplemental Results for details). As described below, Bayesian model comparison provided strong evidence that the selective maintenance model characterized human behavior better than alternatives. In additional simulations, we corroborated that the selective maintenance model was reliably recovered when it generated the behavioral data (exceedance probability [EP] = 1.0), whereas it did not characterize data simulated from other models (EPs < .02; additional details in Supplemental Results).

### Does uncertainty-driven exploration improve foraging success in simulations?

We tested models’ optimality — or foraging success — in multiple novel environments, simulated with a set of complex temporal contingencies with local minima (Supplementary Figure 2). Using the five best parameter sets for each model from an initial search for optimal parameters, we simulated the proportion of possible points earned across 100 novel contingencies. We varied run lengths (40, 60, or 110 trials; detailed in Materials and Methods) to gauge models’ relative performance early and late in learning. The average proportions earned as a function of model and run length are depicted in Figure 4.

**Figure 4.**
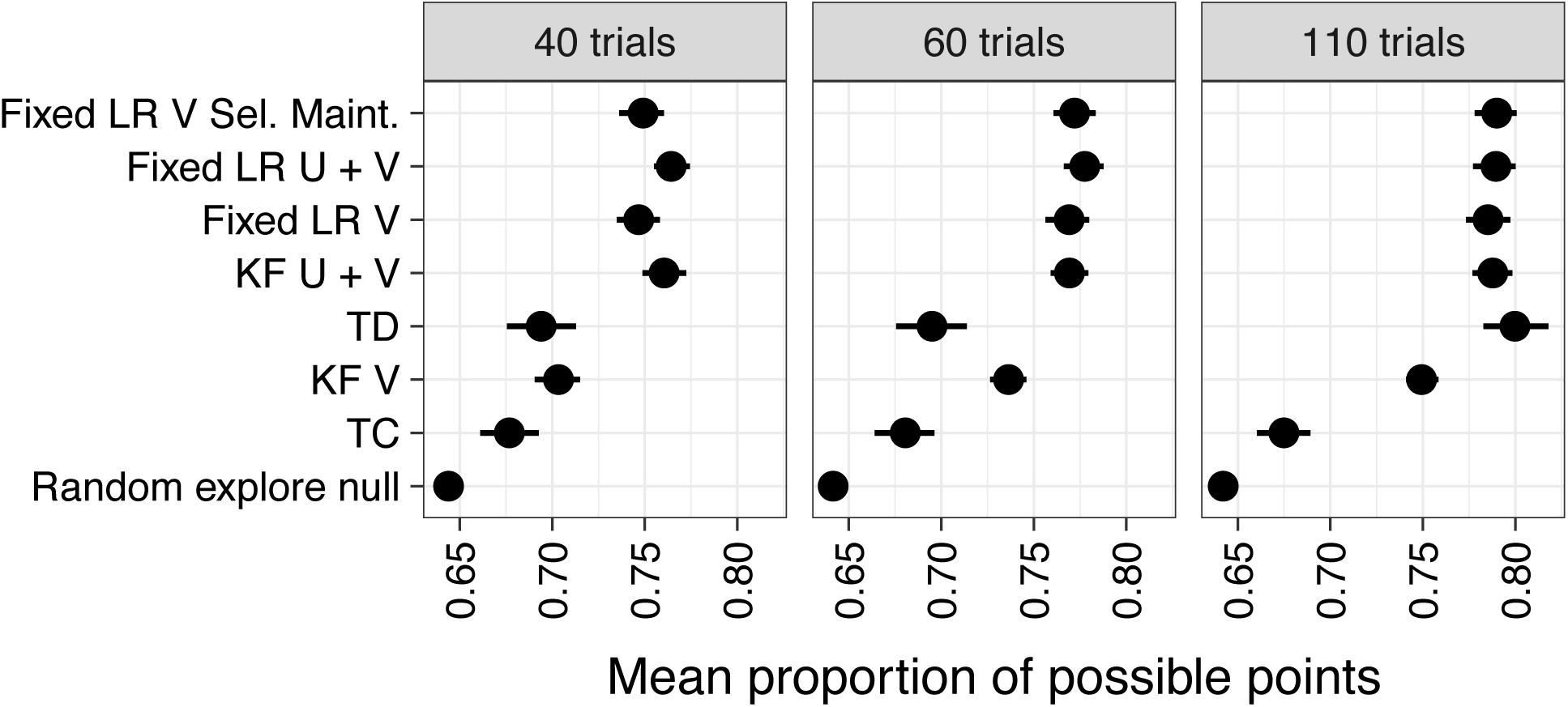
Mean proportion of possible points earned in simulated learning of sinusoidal time-varying contingencies as a function of computational model and run length. Outcomes were drawn from 100 phase-shifted variants of a sinusoidal contingency (details in Materials and Methods). Dots represent the mean, whereas the intersecting line represents the bootstrapped 95% confidence interval around the mean. The model naming scheme is detailed in Table 1. We found significant main effects of model and run length on the proportion of points earned (*p*s < .0001) that were qualified by a model x run length interaction, χ^2^(14) = 228.13, *p* < .0001. Regardless of run length, models that explicitly represented uncertainty (KF U + V and Fixed LR U + V) did not perform significantly better than simpler fixed learning rate models (Fixed LR V Selective Maintenance and Fixed LR V), *adj. p* s > .49. The KF V model performed significantly worse than other SCEPTIC variants at all run lengths, *adj. p* s < .01. TD performed worse than the top four SCEPTIC models at run lengths of 40 and 60 trials (*adj. p* s< .05), but not 110 (*adj. p* > .10). Likewise, TD was not significantly better than TC for 40- and 60-trial runs, but outperformed TC for 110-trial runs, *adj. p* < .001. Finally, TC did not significantly exceed the random exploration null model at any run length, *adj*. *p*s > .10, whereas all other models did, *adj. p*s < .05.

Contrary to our expectation, SCEPTIC variants that leveraged uncertainty to guide choice did not outperform fixed learning rate models guided by value alone, despite having access to information that could be used to sample the action space more systematically. Notably, however, the correlation between the model’s expected value and the underlying contingency was significantly higher early in learning (especially in the first 10 trials) for models that represented uncertainty compared to models that did not, *p* < .0001 (Supplementary Figure 3). Thus, despite yielding a higher fidelity representation of the contingency, uncertainty representation did not enhance overall model performance in simulations.

The selective maintenance model did not perform worse than its full maintenance analog (Fixed LR V; *adj. p* > .10), suggesting that selective maintenance of the value representation did not hamper foraging. Crucially, in a model variant where value traces decayed randomly across the interval, rather than as a function of choice history, performance was impaired (40 trials *adj. p* = .15, 60 trials *adj. p* < 10^-5^; 110 trials *adj. p* = .002; optimized parameter set with γ = 0.24). This demonstrates that selective, but not random, decay promotes adaptive exploitation by maintaining the value of preferred actions (potentially stabilizing a value bump). Finally, to identify boundary conditions where entropy-driven exploration would not suffice, we tested our models in sparse, discontinuous environments with competing value maxima. As detailed in the Supplemental Results, SCEPTIC variants with an explicit uncertainty representation gained a modest advantage. Nevertheless, the selective maintenance model was superior to its full-maintenance equivalent, suggesting that maintaining a subset of valuable actions is efficient even in sparse environments, perhaps in order to avoid the inferior parts of the action space.

### Human Behavior

#### Representation: TBF is superior to TD

SCEPTIC models using TBFs afforded a better fit to behavior than TD (Figure 5a). The representational power of the temporal basis was not simply due to a high number of hidden states (*k* =24, cf. 80 actions tracked by TD). SCEPTIC fits were qualitatively unchanged regardless of the number of basis function elements (data available upon request).

**Figure 5.**
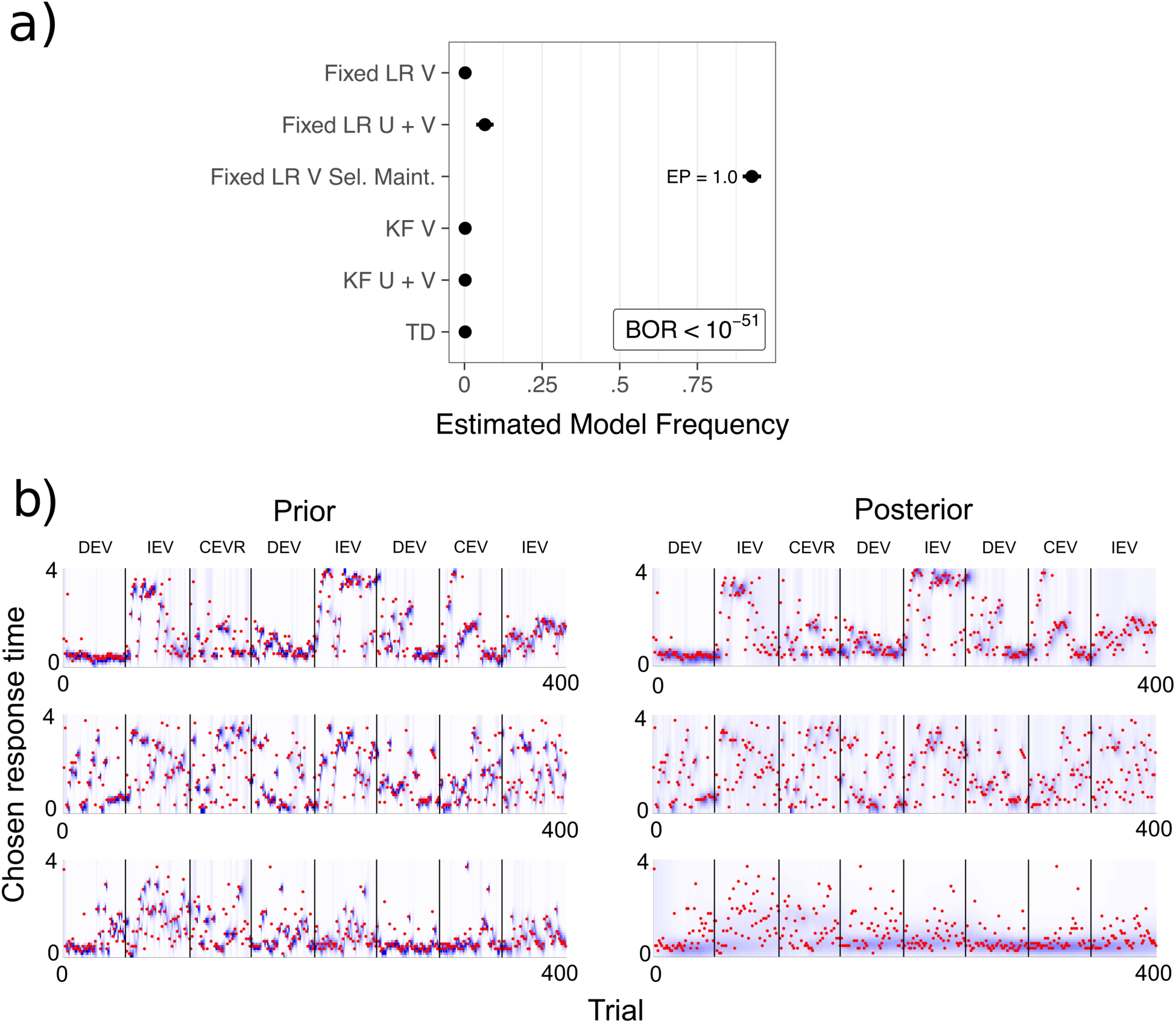
a) Random-effects Bayesian model comparison of SCEPTIC variants and TD. EP = exceedance probability. Dots represent the estimated model frequency, and the intersecting line represents the standard error of the estimate. BOR = Bayesian omnibus risk. Model variants are detailed in Table 1. For a group model comparison including supplemental SCEPTIC variants, see Supplementary Figure 6. b) Trial-by-trial fits of the selective maintenance model in three randomly selected subjects at uninformative prior values of the parameters (left) and at posterior values (right). Subjects’ responses are indicated in red; the model’s posterior predictive density is depicted in blue. Exploitative choices are predicted precisely, whereas exploratory choices marked by large RT swings have lower posterior predictive densities (high entropy). The latter observation is particularly valid at the priors (left), where the temperature of the softmax is fixed at a low value.

#### Selective maintenance of value representations

In a Bayesian model comparison, the selective maintenance model dominated (Figure 5a; EP = 1, Bayesian omnibus risk (29) [BOR] < 10^-51^), indicating that subjects preferred recently visited segments of the interval much more than would be predicted by their long-term expected value. The advantage of the selective maintenance model — measured by the log ratio of free energies vs. all other models — was greater in better-performing subjects, all *r* s(74) > 40, *p*s < 001, except in comparison with TD: *r* (74) = .06, *p* = .48), suggesting that the selective maintenance model captured an adaptive strategy. Moreover, the selective maintenance parameter, γ, correlated significantly with total points earned on the task, *r* (74) = .37, *p* < .001. Posterior predictive checks on the fit of the model to subjects’ behavior suggested that the selective maintenance model captured trial-by-trial variation in behavior, as well as individual differences in response tendencies (Figure 5b).

In addition, we found a significant correlation between γ and nonverbal intelligence, *r* (74) = 0.39, *p* =.0005 (Figure 6) and, more weakly, with verbal intelligence, *r* (74) = 0.23, *p* = .045. Learning rate and performance intelligence were also moderately correlated, *r* (74) = 0.25, *p* = .03. However, when γ, learning rate, and the learning rate x γ interaction were entered into a multiple regression model, they did not predict incremental variance in nonverbal intelligence beyond γ alone, *F* (2, 83) = 1.73, *p* = .19. The relationship between nonverbal intelligence and selective maintenance was not moderated by age (*p* = .19) or sex (*p* = .97).

**Figure 6.**
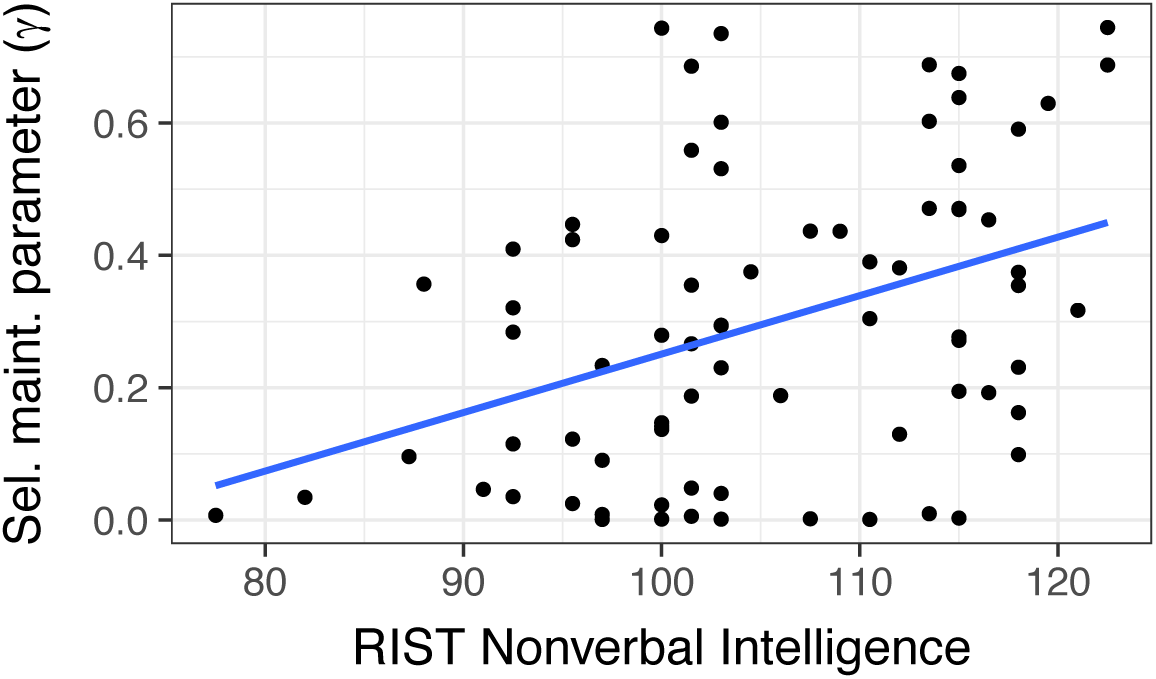
Association between Reynolds Intellectual Screening Test (RIST) nonverbal intelligence score (scaled relative to population norms where *M* = 100 and *SD* = 15) and the selective maintenance parameter. Note that the strength of the association was not substantially weaker when the two lowest intelligence scores were excluded, *r* (73) = .34, *p* =.003.

#### Entropy of the expected value distribution tunes the explore/exploit tradeoff and is shaped by selective maintenance

As illustrated above (Figure 3), selective maintenance of value traces should result in a compressed representation later in learning, as the value of unchosen actions decays toward zero, and entropy decreases. Supporting this prediction, for the selective maintenance model, entropy (information content) was high when participants entered a new contingency, then declined with learning (depicted in Figure 7). Conversely, for the Fixed LR V model, mean entropy was much higher on average (*t* = 246.03, *p* < 10^-16^) and remained relatively stable with learning.

**Figure 7.**
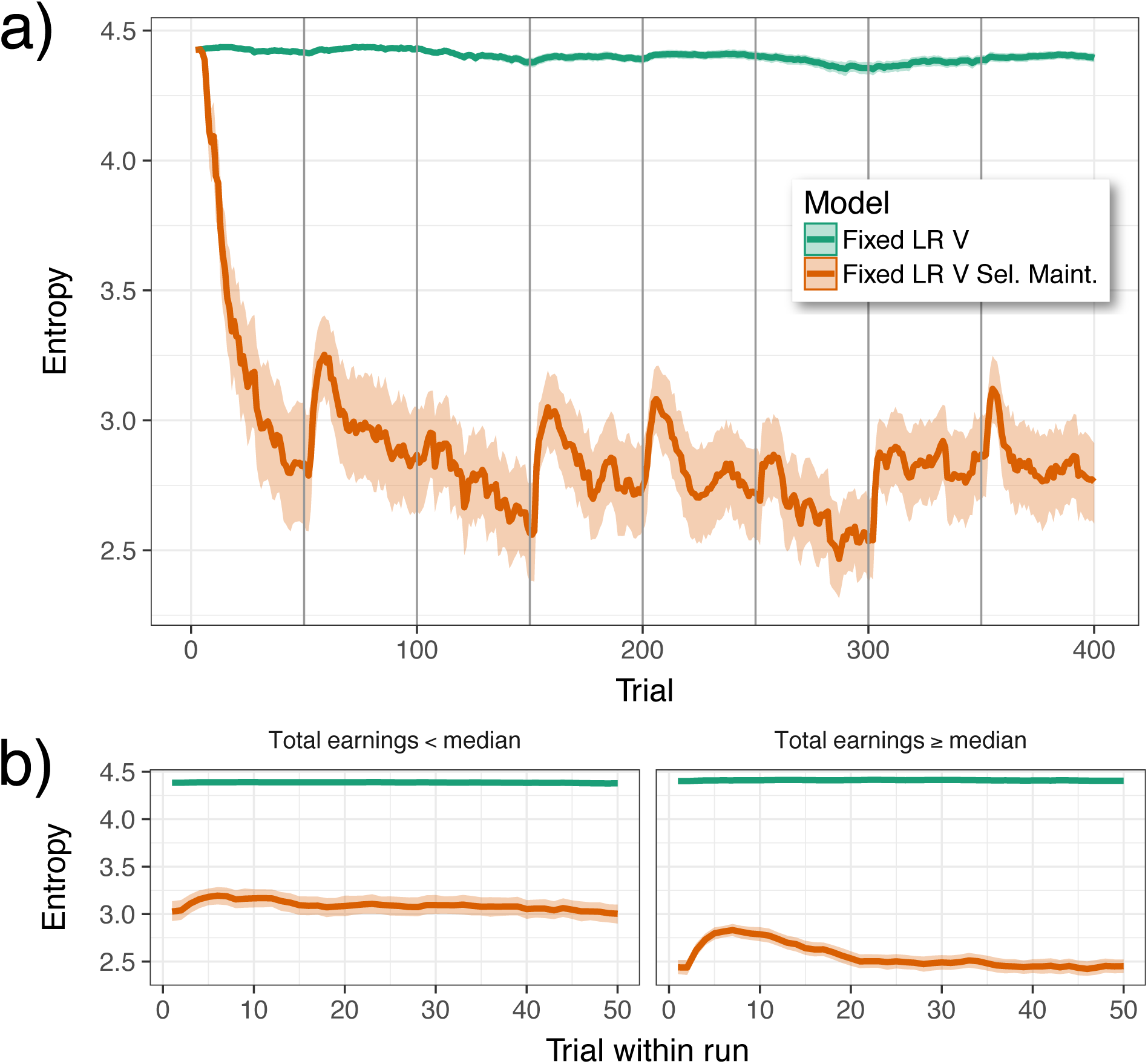
Evolution of value entropy starting from random uniform prior estimates on value. Lines represent the mean entropy across trials, averaging across subjects. Trial-wise entropy was derived from the estimated value distributions of subjects at their best-fitting parameters. Shaded ribbons represent the bootstrapped 95% confidence interval of the mean at each trial. In panel a, dark vertical lines depict boundaries between different contingencies (50 trials each), explicitly signaled to participants. Panel b depicts the average change in entropy, averaging over subjects and runs (excluding run 1); this represents the typical increase in entropy during initial exploration followed by its decline as high-value actions are discovered and exploited. Better-performing subjects (right panel) exhibit proportionately greater entropy increases early in learning under the selective maintenance model, whereas poorer subjects (left panel) have higher mean entropy. Value traces were carried forward from one block to the next, an implementation that resulted in better fits for both models compared to resetting values in each block (data available upon request). Apart from differences in the first few trials of the experiment, the essential dynamics of entropy under the Fixed LR V and Selective Maintenance models are unchanged if the model is initialized with zero prior estimates on value (see Supplementary Figure 7).

Because entropy tunes the exploit/explore tradeoff (30), we hypothesized that early increases in entropy facilitate exploration and the discovery of valuable actions, whereas entropy declines late in learning enable a shift to exploitation, contributing to foraging success. To test this hypothesis, we estimated the effect of entropy early and late in learning (averages in trials 2–10 and 41–50, respectively) on the total number of points earned in each run. As predicted, for the selective maintenance model, higher entropy early in learning predicted greater earnings over the run, *t* = 5.77, *p* < 10^-5^, whereas entropy late in learning was associated with poorer earnings, *t* = −3.85, *p* < .001. Furthermore, we anticipated that greater early:late entropy ratios reflect an adaptive transition from exploration to exploitation and would be associated with better performance. We found strong support for this hypothesis: across subjects, higher early:late entropy ratios were associated with greater total earnings on the task, *r* (74) = .56, *p* < .0001. Crucially, the early:late entropy ratio for the Fixed LR V model (lacking a selective maintenance process), was uncorrelated with performance, *r* (74) = − 16, *p* = .18 (see Figure 8).

**Figure 8.**
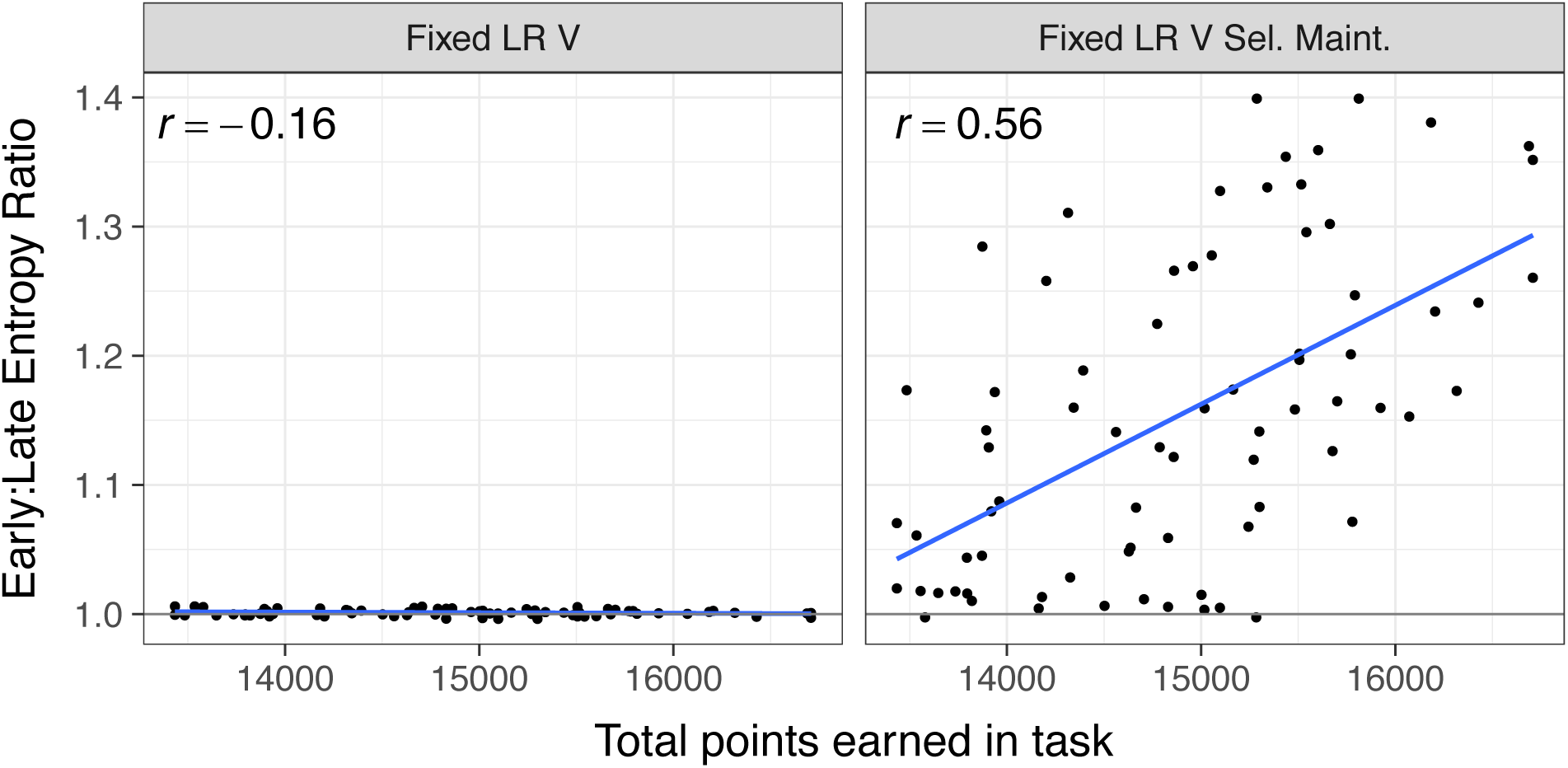
Relationship between early:late entropy ratio and total earnings during the clock task. The early:late entropy ratio was calculated as the quotient of entropy early in learning (trials 2-10) and late in learning (trials 41-50). To account for between-subjects variability in average entropy, for each run, early and late entropy were normalized by the subject’s mean entropy. One extreme high value of the early:late ratio and one participant with low total earnings were Winsorized for plotting and calculations based on regression diagnostics. The correlation between entropy ratio and performance was qualitatively unchanged using the original data: selective maintenance model *r*(74) = 0.53, *p* < .0001. The blue lines denote the least-squares regression line.

The entropy-performance relationship was specific to the selective maintenance model, suggesting that adaptive entropy dynamics are shaped by selective maintenance of value traces. To test this idea, we examined whether the positive association between selective maintenance, γ, and earnings was mediated by the early:late entropy ratio. Corroborating this account, in a path analysis the indirect effect of γ on earnings via the early:late entropy ratio was significant, *β* = .33, *z* = 3.41, *p* < .001 (*p*-values based on bootstrapped standard errors; (31)). Importantly, the direct effect of selective maintenance on performance was nonsignificant (*p* = .80) after accounting for early:late entropy ratio, and the total effect was largely explained by the mediated path, indirect/total effect = .91. Conversely, an alternative model in which individual differences in selective maintenance mediated the relationship between early:late entropy ratios and performance was non-significant, *z* = .26, *p* = .79.

#### Entropy-driven softmax exploration explains RT swings

Our findings are inconsistent with the idea that RT swings (defined as the absolute change in response time on the current trial compared to the previous trial) reflect shifts toward uncertain options. Drawing on the observation that successful learners rely on early entropy to learn the contingency, however, an alternative hypothesis is that RT swings result from high entropy of the expected value representation. Consistent with this account, greater entropy (computed from the selective maintenance model) predicted larger RT swings, *B* = 174.00, *t* = 16.15, *p* < 10^-16^. Importantly, this effect could not be accounted for by many other factors — distance from the point of maximum estimated value, value of the chosen RT relative to the global maximum, distance from the edge of the time interval, trial, the magnitude of the previous RT swing, and whether the previous action resulted in a reward or omission. Even after controlling for these variables and their interactions in a multilevel model, the magnitude of the entropy effect remained essentially unchanged, *B* = 144.04, *t* = 14.19, *p* < 10^-16^(depicted in Supplementary Figure 8). To rule out the possibility that the entropy-exploration relationship reflected high average levels of entropy (e.g., reflecting consistently random responses), rather than trial-level effects, we decomposed entropy into between-run versus within-run variability. Although RT swings were larger in runs with high average entropy (*t* = 16.75, *p* < 10^-16^), high entropy on a given trial (relative to the run mean) predicted larger RT swings on the subsequent trial, and this relationship became stronger as learning unfolded (main effect *t* = 3.26, *p* = .003; trial x entropy interaction *t* = 5.04, *p* < 10^-7^), suggesting a greater role of entropy in continual rather than initial exploration. Interestingly, higher entropy was associated with longer response times, *t* = 10.34, *p* < 10^-16^, consistent with the idea that tracking more value information increases cognitive load, slowing response times or promoting indecision.

#### Uncertainty and exploration

Although the selective maintenance model was uncertainty-insensitive, the next-best model, Fixed U + V, recovered a negative *τ* parameter, indicating uncertainty aversion, for 71 of 76 subjects, *t* (75) = −6.9, *p* < 10^-8^. Within the KF family, the KF U + V model dominated (EP = 1, BOR < 10^-23^) and also recovered negative τ parameter values for 67 of 76 subjects, *t* (75) =-5.8, *p* < 10^-6^. Exploring the large action space of the clock task, participants can shift their response times substantially from trial to trial, particularly early in learning. RT swings were first described by Frank and colleagues (10,11), who viewed them as a form of uncertainty-driven exploration in some individuals. An advantage of the SCEPTIC model is that the tendency of subjects to shift toward or away from the moment of maximal uncertainty (i.e., the RT about which the least is known) can be estimated. To test whether response times were related to uncertainty seeking or aversion, in a multilevel model we regressed trial-wise RT on the previous RT, whether the prior response was rewarded, the RT of maximal value, and the RT of maximal uncertainty. Trial-wise value and uncertainty estimates were obtained from the Fixed U + V model using fitted subject parameters. As expected, there was a strong positive relationship between the highest value option and the chosen response time, *t* = 38.58, *p* < .0001. We also observed a negative association between RTs and the most uncertain option, *t* = −3.09, *p* = .002, indicating that subjects were uncertainty averse. Importantly, the effect of uncertainty was moderated by trial, such that subjects were increasingly averse to the most uncertain option later in learning, RT uncertain x trial *t* = −8.11, *p* < .0001. Finally, consistent with the idea that the U + V model captures individual differences in uncertainty aversion, subjects with more negative τ parameters tended to avoid the most uncertain option to a greater extent, RT uncertain x τ *t* = 3.19, *p* = .001.

#### Are selective maintenance of action values and uncertainty aversion proxies for sticky choice?

Using various implementations of choice autocorrelation (27,32), we ascertained that sticky choice did not account for either value selective maintenance or uncertainty aversion (Supplemental Results).

#### Time-Clock (TC) model

One cannot directly compare fits of the TC model to SCEPTIC and TD because of the different nature of the response variable (a single predicted response time for TC vs. a multinomial choice distribution in SCEPTIC and TD). We did, however, assess the explanatory power of each parameter in the TC model by fitting model variants with an increasing number of parameters, following the order in the model equation (i.e., varying from one to seven parameters; see Materials and Methods). In so doing, we tested whether models that included only descriptive parameters fit substantially worse than models that included RT modulation due to prediction errors, value-based learning, or uncertainty.

Surprisingly, the substantively interesting parameters of TC — expected value of fast versus slow responses (*ρ*), uncertainty-driven exploration (ε), go (α_*G*_), and no-go (α_*N*_) terms — did not contribute substantially to fits. Rather, a comparison among models revealed a strong preference for a model containing three parameters of no interest: mean RT (*K*), choice autocorrelation (λ), and RT of maximum reward (*v*); exceedance probability (EP) = 1.0, Bayesian omnibus risk (BOR) < 10^-35^(Figure 9). In generative simulations using the TC model (based on code from (11), and using published parameters), we replicated the findings of model sensitivity to different contingencies, but observed poor performance in longer learning episodes and under moderate variations in parameters (see Supplementary Figure 9).

**Figure 9.**
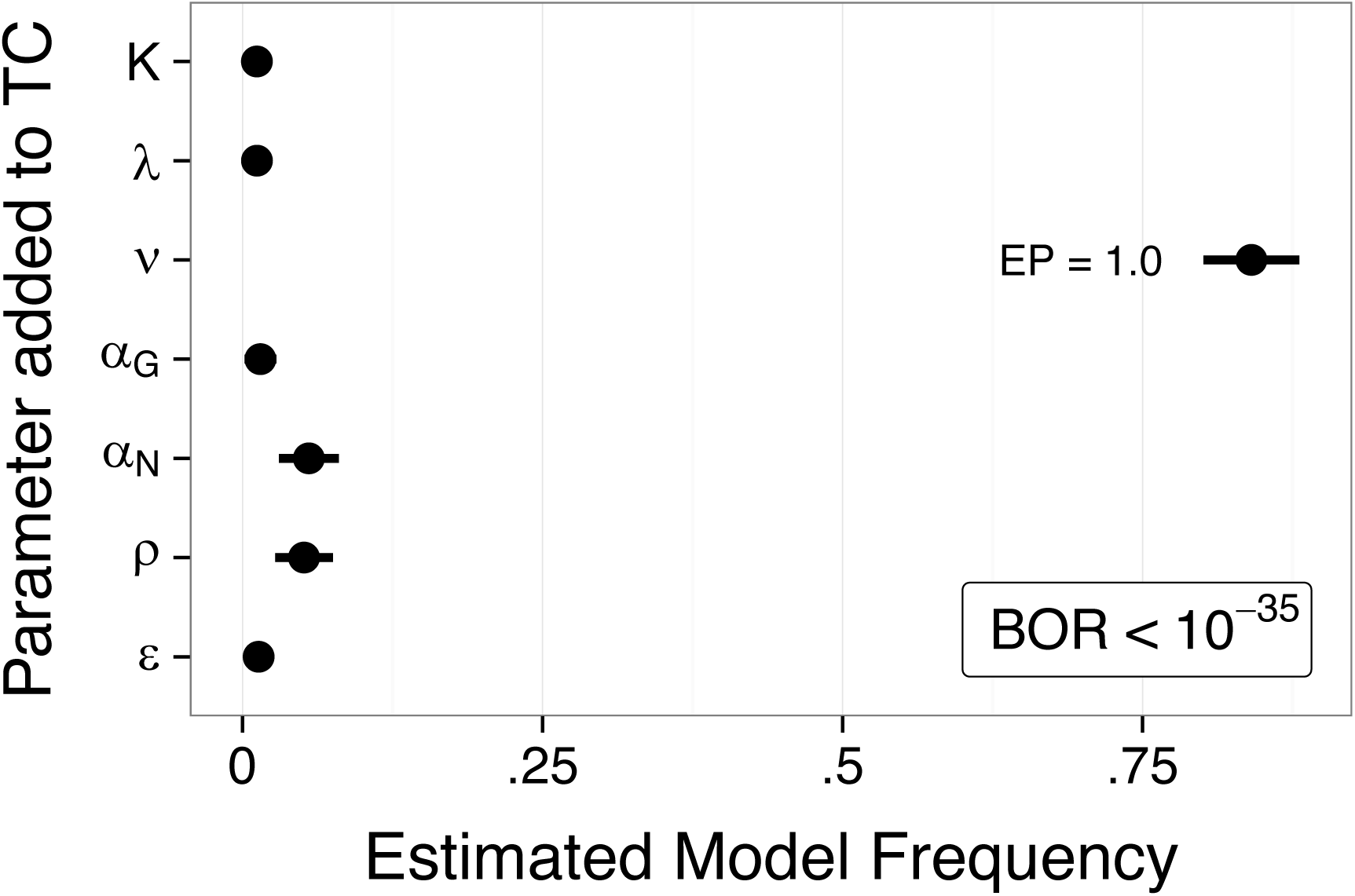
Random-effects Bayesian model comparison of time-clock (TC) model variants incorporating free parameters incrementally (top to bottom). Each tick on the vertical axis represents the addition of that parameter into a TC model variant containing all parameters above it. Thus, models varied from one to seven parameters. EP = exceedance probability. BOR = Bayesian omnibus risk, a measure of statistical risk in group model comparisons quantifying whether chance is likely to explain differences in estimated model frequencies.

## Discussion

Reinforcement-based timing involves exploration of a large continuous action space. We aimed to understand (1) how value information may be represented and maintained in this context, given realistic cognitive constraints, and (2) how information maintenance might shape exploration. Humans’ sampling trajectories reflected selective maintenance: actions with high perceived value were updated by sampling (and prediction error), whereas infrequently sampled action values decayed. Selective maintenance dynamically shaped the information content (entropy) of the model-estimated value representation. Upon entering a new environment, entropy was high during initial exploration and declined later in learning. These dynamics were associated with successful performance and intelligence. This was not the case under full maintenance of value traces, where entropy was high throughout learning and did not predict performance. Consistent with softmax exploration based on a Gibbs/Boltzmann distribution (30), entropy controlled exploration (response time swings). The idea that the shrinking of entropy by selective maintenance facilitates the transition from exploration to exploitation is new to the study of human decision-making.

By contrast, we found no evidence of uncertainty-driven exploration in this context; rather, subjects generally avoided uncertain areas of the time interval to a greater extent than dictated by their reward value. In simulations, uncertainty-driven exploration yielded a more precise representation of the environment early in learning but conferred no appreciable foraging advantage over softmax exploration, with the possible exception of extremely difficult, discontinuous environments where high-value regions were sparse. Extending prior work on TD models of Pavlovian learning (12,13) we found that a neurobiologically plausible temporal basis function representation accounted well for instrumental reinforcement-based timing. Finally, we did not find evidence of computational mechanisms that might control a dynamic learning rate on the clock task, and models with a fixed learning rate afforded the best fit to behavior.

### Information dynamics: selective maintenance of value traces and its effects on entropy

Under the best-fitting selective maintenance model, the information content (entropy) of value traces is high during the initial exploration of a new environment and declines later in learning, facilitating the shift toward exploitation (Figure 7). These information dynamics are more pronounced in successful subjects, indicating that they describe an adaptive approach. Although the idea that adaptive learning involves loss of information may at first seem paradoxical, it fits the intuition that cognitive load is highest when one enters a new (or altered) environment, declining later as one learns to exploit the contingency. Consistent with this account, high entropy was associated with longer response times, which parallel the detrimental effects of working memory load on reinforcement-based timing (33).More generally, the concept of a maximum-entropy information source that emits exploratory actions, as is the case of Boltzmann’s softmax with a uniform input, is not new. It directly relates to Borel’s so-called infinite monkey theorem (34) — proposed as a comment on Boltzmann’s work — where a million monkeys typing randomly eventually produce volumes that will include “books of any nature” (also see Borges’ the Library of Babel; 36).

In simulations, a selective maintenance strategy was generally as successful as comparators that maintained learned values for all sampled actions. In other words, it typically suffices to track a subset of valuable alternatives without maintaining a detailed representation of the rest of the environment. In many individuals, later in learning, the decision function for the selective maintenance model had a peaked unimodal distribution around the perceived value maximum. Epistemically, such dynamics lead one to test a hypothesis about the location of the maximum value (i.e., when in time am I most likely to obtain the best outcome?) rather than tracking the expected value of all options. In this framework, uncertainty about the temporal occurrence of maximum reward is encoded implicitly by the entropy of the value distribution (36).

Moreover, our results suggest one possible computational mechanism of selective maintenance of value representations. Short-term memory traces decay with time, but can be maintained by a refreshing process, such as rehearsal or memory search (17,33,37–39). Extending these observations to reinforcement-based timing, we tested a model where (1) regardless of their valence and magnitude, prediction errors enhance the maintenance of action values and that (2) this enhancement is relatively precise in time following a temporal generalization gradient. Thus, prediction error updates, requiring a retrieval of value traces, can serve as a refreshing process similar to explicit rehearsal or memory search. Similar to the pruning of branching action sequences described by Huys and colleagues (40), TBFs and selective maintenance exemplify heuristic computational mechanisms for reducing the dimensionality of the environment to make learning tractable. Our findings with respect to cognitive constraints on learning also echo Collins and colleagues (14), who found that working memory capacity limits learning in a large action space, accounting for reward learning deficits in schizophrenia. Similarly, Otto and colleagues found that the limits of working memory constrained learning in an environment with a sequential structure (15).

One may note a homology between our algorithmic selective maintenance model and neural network models with attractor dynamics. Basis function elements can be thought of as tuning curves of excitatory neurons, whereas selective maintenance is homologous to inputs from inhibitory neurons. Thus, inhibitory inputs could stabilize the attractor bump in order to maintain a hypothesis about the global value maximum, a testable hypothesis. Reinforcement-based timing is not entirely abolished by any specific brain lesion (41–45), and it is likely that TBF dynamics are found in multiple networks implicated in both timing and reward learning such as the basal ganglia/dopaminergic midbrain, the cerebellum, and the premotor cortex (46–48). It is also likely that selective maintenance is an active process mediated by circuits involved in time-based resource allocation, such as the frontoparietal networks. We predict that entropy should scale with the number of active value-sensitive elements in these circuits.

### Uncertainty-Driven or Entropy-Driven Exploration?

Intuitively, when choosing among a multitude of actions with uncertain reward values, an agent should benefit from uncertainty-driven exploration, at least early in learning (10,49). Yet the resulting need to appraise uncertainty complicates the already intractable exploration/exploitation dilemma (50), giving rise to a number of specific problems and objections. Our simulations showed that optimizing a single tradeoff between directed, uncertainty-driven exploration and exploitation, despite yielding a more precise map of the environment early in learning, did not confer an advantage over models with undirected, stochastic exploration. These results likely generalize beyond temporal contingencies to other action spaces with correlated returns along a continuous dimension and may help to explain why humans do not engage in uncertainty-driven exploration in other tasks (7,51). Importantly, cognitive constraints likely limit tracking of uncertainty. If we assume that uncertainty representation decays like other memory traces, representations in the infrequently visited (and most uncertain) regions of the action space would be most subject to decay, causing the uncertainty estimate to revert toward a prior (cf. 51) and hindering directed exploration. The total loss of information about unsampled actions given a fixed representational capacity and a decay of uncertainty estimates also scales with the size of the action space.

Although uncertainty-driven exploration can be efficient initially in complex environments (e.g., large spatial landscapes (23) or discontinuous environments with sparse high-value regions), the tasks used in studies of human value-based decision-making typically emphasize continual exploration, where one exploits the perceived contingency and periodically explores to adjust the policy to the environment. Our results suggest that an implicit representation of uncertainty expressed in the entropy of action probability distribution may be necessary for continual exploration. That said, things change in sparse, discontinuous environments in which only a small fraction of possible actions are reinforced. In additional simulations (Supplementary Figure 4 and 5) of such environments, models incorporating uncertainty-guided exploration gained a small advantage over value-guided choice alone. At the same time, these very challenging environments could not be learned by TD probably because of their temporal discontinuity, which defeats value back-propagation. Considering that and the human performance on the much simpler monotonic contingencies (Figure 1), we doubt that an average person can learn sparse, discontinuous contingencies reliably even given hundreds of trials.

### Basis Function Representation of a Temporal Contingency

The SCEPTIC model represented expected value using several radial basis functions with contiguous and overlapping temporal receptive fields. We note that discrete Gaussian elements were employed here for computational convenience, and we make no claims about the superiority of this solution over alternatives such as cosine basis (52). Extending earlier findings in Pavlovian conditioning (12,13), we found that TBF representations afforded a better fit to human instrumental behavior on the clock task than a delay line TD model. TD has been previously reported to fail on the clock task due to erroneous back-propagation of value from later to earlier points in the interval (24). Importantly, our TD model with a task-specific state space partition successfully overcame this problem, as indicated by its foraging success. We can thus be confident that SCEPTIC’s superiority is specifically due to the combination of TBF value representation and the strategic consideration of the entire interval in the choice rule. Like Ludvig, Sutton, and Kehoe (12,13), we did not attempt to estimate the functional form, number, and placement of TBF elements, fixing them at reasonable priors. New experiments are needed to constrain the parameterization of TBF with behavioral and physiological data. One testable hypothesis — articulated but not tested by Ludvig and colleagues — is that the density of elements decreases progressively with the passage of time within the interval, which would reflect Weber’s law (53). This question relates to the broader unresolved problem of interval adaptation (47,53), or in computational terms, how the agent learns to optimally place the TBF elements based on time intervals experienced in a particular context.

### Limitations

While our data support entropy-driven continual exploration on the clock task, they do not rule out the possibility that initial exploration is driven by uncertainty seeking. That said, we tested a model variant (KF U → V) designed to switch from initial exploration to later exploitation, which was inferior in model comparisons (details in Supplemental Materials). The time course of exploration (Figure 1) and information dynamics (Figure 7) suggest that the period of initial exploration on the clock task may be as short as 5-8 trials. One possibility is that humans track uncertainty explicitly during the first few trials in order to seed potentially valuable actions in the softmax function, essentially defining a subset of eligible actions for further refinement in continual exploration. Further, in environments where participants are explicitly told that they will repeatedly sample a stable contingency (i.e., there is a longer horizon for learning), they are more likely to use directed exploration to identify the best action than when they have only a single choice (54). More speculatively, the cognitive load of an explicit uncertainty representation combined with a high-entropy value representation may be overwhelming early in learning. On the other hand, the shift toward continual exploration using a Boltzmann strategy may ease working memory demands. Our null findings with respect to learning rates should be taken with caution since there are a number of other approaches to volatility-modulated learning rates that could be potentially adapted to SCEPTIC (51,55,56). Finally, our analyses did not address the issues of opportunity cost and the intrinsic cost of waiting (52), important independent influences on behavior, which will need to be examined in the context of the clock task.

## Conclusions

In contrast to previous proposals for resolving the exploration/exploitation dilemma at policy level, we show that this can be also accomplished at the level of updating and maintaining the value representation. In this study of reinforcement-based timing a simple selective value maintenance strategy reduced information load and facilitated the transition to exploitation, further work is needed to evaluate alternative information maintenance strategies and their putative neural implementations. At the same time, our findings broadly align with emerging evidence that the implementation of optimal inference is limited by representational capacity (2) and that heuristic approaches to reinforcement learning are more likely to be effective in complex environments (20,57).

## Materials and Methods

### Ethics Statement

Participants and/or their legal guardians provided informed consent or assent prior to participation in this study. Experimental procedures for this study complied with Code of Ethics of the World Medical Association (1964 Declaration of Helsinki) and the Institutional Review Board at the University of Pittsburgh (protocol PRO10090478). Participants were compensated $75 for completing the experiment.

### Participants

We enrolled 76 normally developing youth and young adults, aged 14 to 30 (*M* = 21.32, *SD* = 5.10). Thirty-nine participants (51.3%) were female. Prior to enrollment, participants were interviewed to verify that they had no history of neurological disorder, brain injury, pervasive developmental disorder, or psychiatric disorder (in self or first-degree relatives).

### Procedure

Participants completed eight runs of a reinforcement-based timing task (hereafter called the “clock task”) during an fMRI scan in a Siemens Tim Trio 3T scanner. Runs consisted of fifty trials in which a darkened circle resembling a clock hand revolved 360° around a central stimulus over the course of four seconds (Figure 1a). Participants pressed a button on a button glove to end the trial and receive a probabilistic reward. Time-varying contingencies were taken from the paradigm developed by Moustafa and colleagues (24) and included monotonically increasing expected value (IEV; reinforcing late responses), decreasing expected value (DEV; reinforcing early responses), constant expected value (CEV), and constant expected value–reversed (CEVR; see Figure 1b). The CEV and CEVR conditions had constant expected value for all response times, but varied in probability and magnitude. The central stimulus was a face with a happy expression or fearful expression, or a phase-scrambled version of face images intended to produce an abstract visual stimulus with equal luminance and coloration. Faces were selected from the NimStim database (58). All four contingencies were collected with scrambled images, whereas only IEV and DEV were also collected with happy and fearful faces. The emotion manipulation and fMRI results will be reported in separate manuscripts because they are not central for the validation of our model.

Participants also completed the Reynolds Intellectual Screening Test (RIST), a brief inventory of verbal and nonverbal intelligence (59) consisting of a verbal subtest measuring verbal reasoning and vocabulary, as well as a nonverbal subtest in which examinees identify which stimulus does not belong with the others in a series of progressively more abstract displays. The RIST has strong test-retest reliability and convergent validity, and correlates highly with full assessments of intellectual ability. The RIST was administered by personnel proficient in psychological testing and supervised by one of the authors (MNH). In our sample, the average RIST Index (a measure of overall intellectual ability) was 105.46 (*SD* = 9.39; range = 80–129).

### StrategiC ExPloration/ExPloitation of Temporal Instrumental Contingencies (SCEPTIC)

#### Temporal basis representation

The SCEPTIC model represents time using a set of unnormalized Gaussian radial basis functions (RBF) spaced evenly over an interval *T* in which each function has a temporal receptive field with a mean and variance defining its point of maximal sensitivity and the range of times to which it is sensitive, respectively (a conceptual depiction of the model is provided in Figure 2). The number, width, and spacing of these basis functions can be varied without loss of generality to more substantive parts of the reinforcement learning model, although a richer basis set can represent more fine-grained temporal information. A set of overlapping radial basis functions (RBFs) provides an efficient approximation of an arbitrary function, *f* (*T*), over a finite interval (60,61). From a biological standpoint, the advantages of this approach are that 1) given a fixed number of basis functions,approximation imprecision scales with the length of the interval (53); and 2) it is compatible with accounts of response-sensitive neurons with distinct temporal tuning, for example in the medial premotor cortex (62).

The primary quantity tracked by the basis is the expected value of a given choice. To represent time-varying value, the heights of each basis function are scaled according to a set of weights, W=.[*w*_1_,. *w*_2_, … *w*_*b*_]. The contribution of each basis function to the integrated value representation at a particular moment in time, *t*, is represented by multiplying the relevant weight by the *b* th RBF:

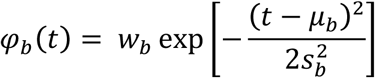

And more generally, the expected value function on a trial *i* is obtained by the evaluation of the basis across time:

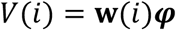

#### Parameterization of temporal basis functions in SCEPTIC

In order to represent temporal decision-making during the clock task, where the probability and magnitude of reward varied over the course of four-second trials, we spaced the centers of 24 Gaussian RBFs evenly across the discrete interval and chose a fixed width, 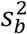, to represent the temporal variance (width) of each basis function. Because of the challenges of approximating functions over discrete intervals (the Runge phenomenon; see (60,63)), two additional modifications of the basis were required to obtain an accurate function approximation: (1) the centers of some basis functions were extended beyond the interval to be represented, and (2) weight updates were performed using a truncated Gaussian basis such that the area under the curve within the time interval of interest was equal for all basis functions (additional details provided in Supplementary Methods).

Although it is plausible that a system coding time-dependent information about rewards may update its temporal width, height, or center of maximal sensitivity on the basis of reinforcement, here we updated only the heights of each basis function based on the reinforcement history. For treatments of basis function adaptation in reinforcement learning models, see (64) and (65). For the widths of the RBFs, 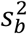, we chose a moderate degree of overlap between adjacent basis functions in order to provide reasonable coverage of each moment within the time interval. More specifically, 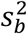 was chosen such that the distribution of adjacent RBFs overlapped by approximately 50%, but overlap between 30% and 70% provided similar results. The parameterization of the temporal basis is not a crucial component of the SCEPTIC model, and another temporal basis (e.g., piecewise polynomial splines or discrete cosine transform) would likely yield similar results.

#### Updating expected value on the basis of reinforcement

A straightforward model of temporal instrumental learning can be specified by combining the delta learning rule (66) with the temporal representational structure defined above. More specifically, the weight for a basis function *b* can be updated according to the equation:

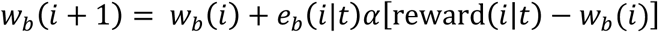

where *i* is the current trial in the task, *t* is the observed response time, and reward(*i* |*t*) is the reward obtained on trial *i* given the choice *t*. The effect of prediction error is scaled according to the learning rate *α* and the temporal generalization function *e*_*b*_. Of note, this learning rule updates the weight of each basis function, *w*_*b*_(*i*), individually without assuming knowledge of the integrated value representation, *V* (*i*). Thus, the value function approximation does not converge absolutely on the temporal distribution of rewards, instead preserving relative differences (i.e., the rank ordering) of alternative values. We note that to make the model converge on the underlying value function, the learning rule of SCEPTIC variants can be altered to compute prediction errors as the difference between actual reward and the integrated value representation, K(L). Implementing such a function-wise update does not qualitatively change any of our substantive results. This alternative, however, suffers from lower identifiability and provides a poorer fit to subjects’ behavior. It also requires a stronger assumption about value representation, namely that basis functions query each other during learning to update their weights (details available from authors upon request). We acknowledge that Ludvig and colleagues’ TBF model of Pavlovian learning applies function-wise value updates (12), and the reasons for the apparent superiority of elementwise updates here remain to be investigated.

To take advantage of temporal generalization, it is crucial that feedback obtained at a given response time *t* is propagated to adjacent times to avoid tracking separate value estimates for each possible moment. Thus, to represent temporal generalization of expected value updates, we used a Gaussian RBF centered on the response time *t,* having width 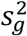.and normalized to have an area under the curve of unity. The eligibility of a basis function φ_*b*_.to be updated by prediction error is defined by the area under the curve of its product with the temporal generalization function:

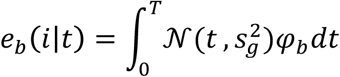

This parameterization leads to a scalar value for each RBF between zero and one representing the proportion of overlap between the temporal generalization function and the receptive field of the RBF. In the case of perfect overlap, where the response time is perfectly centered on a given basis function and the width of the generalization function matches the basis 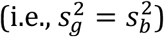 *e*_*b*_. will reach unity, resulting a maximal weight update according to the learning rule above. Conversely, if there is no overlap between an RBF and the temporal generalization function *e*_*b*_ will be zero and no learning will occur in the receptive field of that RBF.

#### Choice rule

Finally, having defined a framework for tracking and updating estimates of expected value over time, SCEPTIC variants select an action based on a softmax choice rule, analogous to simpler reinforcement learning problems (e.g., two-armed bandit tasks (67)). For computational speed, we arbitrarily discretized the interval into 10ms time bins such that the agent selected among 500 potential responses. The agent chose responses in proportion to their expected value:

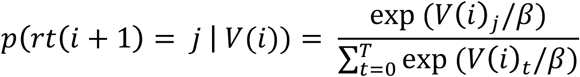

where *j* is a specific response time and the temperature parameter, *β*, controls the sharpness of the decision function (at higher values, actions become more similar in selection probability).

The softmax function has two potentially desirable properties in the temporal instrumental learning context. First, if several actions are associated with similar expected value, even if substantially separated in time, they will be selected with similar probability. Second, by virtue of the temporal basis representation (where reinforcement information is generalized in time), response times adjacent to the global maximum of learned expected value are more likely to be selected, promoting temporally local exploration of advantageous areas. To highlight the advantages of the softmax policy, we contrasted it with a γ-greedy choice rule, which was predictably inferior in optimality simulations (details provided in Supplemental Materials).

#### Overcoming cognitive constraint on value representation: the selective maintenance model

We tested a selective maintenance model (Figure 3) in which basis weights reverted toward zero in inverse proportion to the temporal generalization function:

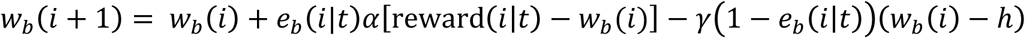

where γ is a selective maintenance parameter between zero and one that scales the degree of reversion toward a point *h*, which is taken to be zero here, but could be replaced with an alternative, such as a prior expectation.

### Representing value and uncertainty according to the Kalman filter

The Kalman filter (KF) is a classic Bayesian approach to estimating the expectation (mean) and uncertainty (variance) of a Gaussian process that unfolds in discrete time (for a classic example of Kalman filters in reinforcement learning models, see (68)). We also note that Frank and colleagues tested a Kalman filter variant of their model to track expected value, rather than the probability of prediction error, and our work built on this useful insight. In the SCEPTIC model, each basis function can be reconceptualized as a Kalman filter that tracks information about both the expected value of a response (i.e., the mean) in its temporal receptive field as well as uncertainty about expected value. Crucially, integrating across basis functions, KF variants of the SCEPTIC model represent both time-varying value and uncertainty functions (*V* and *U*, respectively), enabling policies that integrate information from both sources.

Compared to SCEPTIC variants described above that rely on a fixed learning rate to update basis weights, there are three major differences for KF variants: 1) the effective learning rate (gain) declines with experience such that early outcomes have the greatest effect on learning, 2) the model tracks the evolution of uncertainty about expected value, and 3) the choice rule (policy) for some models involves a tradeoff between exploratory and exploitative influences. The learning rule for KF SCEPTIC variants is:

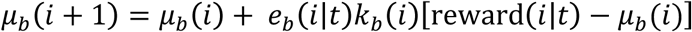

where μ_*b*_.L.represents the contribution of basis function *b* on trial *i* to the expected value function, *V* (*i*). The gain (learning rate) for a given basis function on trial *i* is defined as

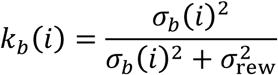

where 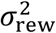 represents the expected volatility (measurement noise) of the environment. Here, we provided the model the variance of returns from a typical run of the experiment as an initial estimate of measurement noise, although other priors led to similar model performance. We also initialized prior estimates of uncertainty for each basis function to be equal to the measurement noise, 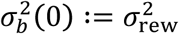, leading to a gain of 0.5 on the first trial (as in (69)).

Under the KF, the contribution of each basis function to uncertainty about expected value is represented as the standard deviation of its Gaussian distribution. Likewise, posterior estimates of uncertainty about responses proximate to the basis function *b* decline in inverse proportion to the gain according to the following update rule:

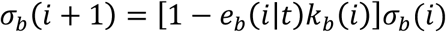

Note that the temporal generalization function *e*_*b*_(*i* |*t*).is parameterized identically across SCEPTIC variants and is used in KF variants to update both value and uncertainty estimates. Extending the temporal representation described above, for KF variants, estimates of the time-varying value and uncertainty functions are provided by the evaluation of the basis over time:

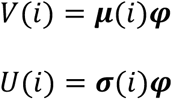

#### Integrating uncertainty and expected value in response selection under KF SCEPTIC variants

Early in learning, expected value will be low for most responses and uncertainty will be high, whereas the converse will be true late in learning. Thus, a policy that combines value and uncertainty may confer particular advantages because uncertainty-driven responses early in learning would facilitate more robust and efficient sampling. The KF U + V policy represents a decision function, *Q*. (*i*)., as a weighted sum of the value and uncertainty functions according to a free parameter, *τ*. As uncertainty decreases with sampling and expected value increases with learning, value-related information will begin to dominate over uncertainty. Positive values of *τ*.promote uncertainty-driven exploration, whereas negative values yield uncertainty aversion.

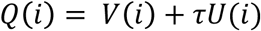

To ensure that our model comparison results were robust to the specific implementation of the uncertainty-sensitive choice rule and dynamic learning rate, we tested a number of alternatives, detailed in the Supplemental Material, all of which proved to be inferior to the variants described above.

#### Sensitivity analyses: choice autocorrelation (sticky choice)

To rule out the possibility that the Selective maintenance model fit well because it better represented sticky choices, we extended SCEPTIC models with two choice autocorrelation functions (ACF): a simple first-order autoregressive (AR[1]) ACF and an ACF extended over multiple trials (32). The AR(1) model modulated the probability of choosing a given time point *t* in a trial by *χ · π* ^|*t –rt* (*i*)|^ (we omit the trial index *i* for simplicity in this paragraph), where. *χ* is the autocorrelation parameter and π is the temporal generalization parameter, followed by divisive normalization. The extended ACF maintained for each time point *t* an index *c*_*t*_ of how recently it was chosen. When *t* was chosen, *c*_*t*_ was set to 1; otherwise it decayed by a factor λ. The probability of *t* being chosen in the softmax was a function of its value and the additive term *χ* ·*c*_t_. We also tested a version of extended ACF with temporal generalization implemented using Gaussian smoothing with a kernel representing a temporal generalization parameter, but this resulting model had inferior fits (data available upon request).

### TD (*Q*-learning) Model

To design a robust benchmark for the clock task, we amended the standard *Q*-learning model in two ways. First, to overcome the problem of erroneous value back-propagation, where earlier responses become over-valued (24), we set up the state space such that each time step *t* has a pair of actions, *A* ={*wait, respond*}. For wait actions, we ensured that value was appropriately back-propagated only to previous waits:

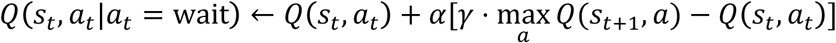

where *α* is the learning rate and *γ* is the discount parameter. We use the conventional *Q* (*s*_*t*_, *a*_*t*_)___.notation here, but since time steps *t* always map onto the same states (potential response times), *s*_*t*_ is redundant and *Q* (*a*_*t*_) would suffice. Conversely, because *respond* actions led to the absorbing terminal state, ending the trial, they were directly updated by actual rewards and not by back-propagation:

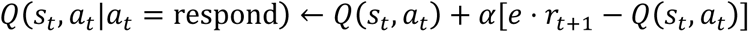

where *e* is the eligibility trace or credit for a reward assigned to a given action based on its temporal proximity. We assumed *e* to decay exponentially over time.

Second, we employed a modified *ε*-greedy choice rule to ensure that the agent explored the later part of the interval:

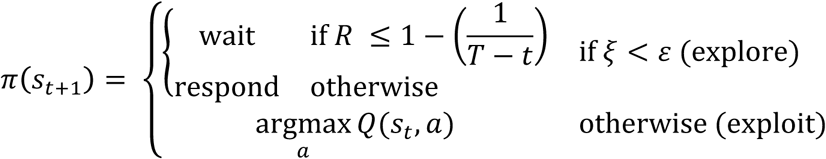

where ε is the exploration/exploitation parameter, *t* is the current time step, *T* is the total number of time steps, and ξ and *R* are [0,1] uniform random numbers drawn at each time step. While a simple *ε*- greedy agent does not effectively explore later time steps because of the fixed exploration probability, the 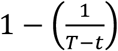 term produces more exploratory *waits* early in the trial and more *responses* late in the trial. A SARSA model was also tested, but is not included here because it was inferior to TD in almost every respect (data available upon request).

### Frank Time-Clock (TC) Model

To date, the TC model of Frank and colleagues (11) has been applied in behavioral, genetic, and neuroimaging studies of uncertainty-driven exploration and dopaminergic influences on response times (10,24). The TC model represents trial-wise response times on the clock task as a linear combination of several potentially neurobiological processes:

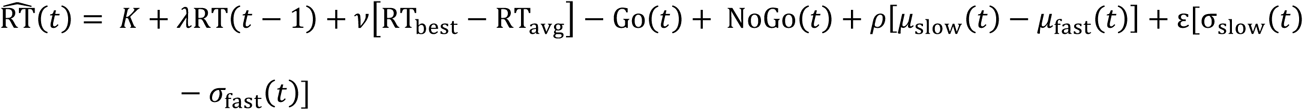

The details of each parameter and the underlying representation are provided in the Supplemental Methods. With respect to value-based decisions, the TC model separately updates the probability of a positive prediction error (PPE) for RTs that are slower or faster than the subject’s average (*μ*_slow_ and *μ*_fast_, respectively). With learning, the model predicts that subjects shift toward faster or slower RTs that are associated with a greater expectation of a PPE according to a free parameter, *ρ*.

### Data Analysis

#### Tests of model optimality

In order to test and compare the efficacy of each model in learning temporal contingencies, we identified parameter sets that maximized the quality of choices in simulations of the clock task. More specifically, for each model, we fit parameters using a genetic algorithm (*ga* function in MATLAB) that returned the greatest summed expected value across 60 runs consisting of 50 trials each. The temporal contingency was based on two sinusoidal functions for expected value and probability (see Supplementary Methods and Supplementary Figure 2 for details). Next, we validated the robustness of parameter sets to contingencies that were not part of the parameter optimization. For this step, we simulated rewards earned by each model at the five best parameter sets from optimization. Each model was exposed to 100 instances of a randomly phase-shifted variant of the sinusoidal contingency and made choices according to its parameters. We then analyzed the distribution of returns for each model using multilevel regression models (*lmer* function in *R* 3.2.5) where contingency instances were modeled by a random intercept (since contingencies were drawn from the population of possible contingencies) and model was treated as a fixed factor. Because models differed considerably in the variability of earnings across replications, we allowed for heteroscedastic residual variance by model.

Because the objective functions for most models were non-convex and prone to local minima, optimization using a genetic algorithm (*ga* function in MATLAB) was repeated 100 times using different random starting values spanning the parameter space for the initial population. This resulted in a distribution of returns on policy, as well as multivariate distributions of parameters for each optimization. These data provided information about the ability of each model to learn the temporal contingency when parameters were tuned for that environment.

To compare the optimality of different models in solving temporal instrumental contingencies, we estimated a multilevel model in which performance was regressed on model and run length (40, 60, or 110 trials); parameter set (best 5 sets for each model) and replication (100) were treated as random effects. Simple effects tests of model for each run length were estimated adjusted *p*-values were computed according to the multivariate distribution of coefficients to maintain a familywise error rate of .05 for each run length (70). We varied run lengths between 20 and 200 in increments of 5, but chose to report a subset of contingencies that were illustrative model performance changes as a function of run length.

#### Identifiability

In addition to comparing the ability of each model to solve temporal instrumental problems in simulated environments, we tested their recovery of each parameter in simulated data. As in conventional simulation studies of estimators (for a useful treatment in a neuroscience context, see (71)),we focused specifically on variance (i.e., dispersion of estimated values around the population value, reported as *R* ^2^ between original and recovered parameters) and parameter bias (i.e., systematic deviations between population and estimated values). For each model, we simulated the behavior of 100 agents by drawing all parameters from the uniform distribution and fitting all parameters at once. For models using a softmax choice rule, the temperature parameter was not recoverable. Since this parameter tends to absorb misfit and account for exogenous factors when fitting behavior, we fixed it here at 0.1, a value corresponding to high exploitation that highlighted more substantive parts of the model. To avoid bias, we took a similar approach when fitting TD. We estimated best-fitting parameters using the *ga* in MATLAB similarly to our optimality tests.

#### Subject behavior fitting

We fit computational models to participant behavior in a deterministic state-space framework using a variational Bayes approach (VBA) implemented in MATLAB (72). An important advantage of this approach is that the relative evidence for different models can be compared using random effects Bayesian model comparison (BMC) (73). VBA parameterizes the choice history in a state-space framework consisting of dependent variables (i.e., time series to be predicted by the model), hidden states (i.e., latent quantities to be tracked over trials), and evolution and observation functions that define the dynamics of hidden state transitions and the model-predicted output, respectively (for details, see (72)). For SCEPTIC models, the hidden state vector was composed of basis function weights representing expected value, which were initialized from a random uniform prior distribution spanning the range of values on the underlying contingencies. For variants that also tracked uncertainty, posterior uncertainty estimates were also tracked for each basis function and initialized as the variance of values across all timesteps in the distribution. We used uninformative Gaussian priors (*M* = 0, *SD* = 10) for all other free parameters.

For *Q*-learning, the hidden state vector consisted of estimated *Q* values for respond and wait actions at each time step (80 total hidden states). Similar to SCEPTIC, we used uninformative Gaussian priors (*M* = 0, *SD* = 10) for both free parameters.

For TC, the hidden states tracked by the model were a) the response time associated with the largest reward experienced in a block, b) the *α* and *β* hyperparameters for two beta distributions tracking the value and uncertainty of slow and fast responses, c) the expected value of each choice tracked according to a delta-rule model with learning rate of 0.1, and d) the value of Go and NoGo decision signals, e) the locally averaged response time, RT_locavg_. Model parameters were initialized with broad uninformative priors, although because of differences in scaling and parameterization (e.g., learning rates vary between zero and one), the distributional form of parameters varied (see Supplemental Table 1). Initial values for parameters were chosen based on Frank and colleagues (11) and code generously provided by Michael Frank. Parameter distributions were chosen based on optimization bounds in the previous TC implementation, as well as observations about TC parameters in this and previous datasets.

*Note*. The codes and data for all analyses reported in this paper are publicly available at: https://github.com/DecisionNeurosciencePsychopathology/temporal_instrumental_agent

## Acknowledgements

The authors thank Michael J. Frank for helpful comments on validation and comparison of reinforcement learning models, as well as codes for the experimental paradigm and time-clock model. We thank Jonathan Wilson for help with figure preparation and model implementation. We are grateful to Peter Molenaar for useful suggestions regarding adaptive filtering and autoregressive processes. We appreciate Yael Niv’s comment about the role of basis representation in our learning rule. We also thank Michael Woodford for his suggestion of examining information theory measures in behavioral analyses. We thank Beatriz Luna for help with data collection and quality assurance.

This research was performed, in part, using resources and the computing assistance of the Pennsylvania State University Institute for CyberScience Advanced CyberInfrastructure (ICS-ACI).

## Supplementary Figure Captions

*Supplementary Figure 1.* Identifiability of model parameters in simulations. Original parameters used in simulations (x-axis) vs. recovered parameters (y-axis). Parameters for all models in the SCEPTIC fixed learning rate family (left column) were recovered with high precision and minimal bias. Among the SCEPTIC KF models (middle column), KF process noise and KF U+V parameters were recovered reliably. Only some parameters in the KF U → V and KF volatility model were identified. None of the TC parameters (right column, top) were identified. In TD, *ε* was identified while *α* was not.

*Supplementary Figure 2.* Expected value (EV), probability, and magnitude of rewards for sinusoidal temporal contingency used for model optimality comparisons.

*Supplementary Figure 3.* Median correlation between trial-wise model-estimated value and true value across 500 simulated contingencies as a function of SCEPTIC parameterization. Shaded ribbons represent the standard error of the median estimated using a LOESS smoother. Contrary to our prediction, models that allowed for uncertainty-driven exploration (especially KF U → V and KF U + V) did not have an advantage over similar fixed learning rate models in optimality tests. We expected that including uncertainty in the SCEPTIC choice rule would confer an advantage early in learning because the agent would recover a higher fidelity representation of expected value across the entire action space. Compared to a simple softmax choice rule over the expected value vector, V(i), uncertainty-driven sampling is more likely to sample the action space systematically and develop a better representation of the contingency. This advantage should be especially pronounced early in learning because uncertainty-driven sampling enhances the unique information gained by each action. More specifically, incremental information is maximized by choosing the most uncertain action each time (an extension of entropy reduction; [1]).

To test the hypothesis that models that included uncertainty in the choice rule would more rapidly recover an approximation of the true reinforcement contingency, for each model we computed the Pearson correlation between the trial-wise estimate of expected value, V(i), and the true reinforcement schedule. This step was repeated for each of the 500 simulated replications/contingencies, generating a distribution of correlation estimates at each trial. The tradeoff between exploration and exploitation means that an agent that explored until it had very little uncertainty about each action would do quite poorly on the clock task (unless there were an inordinate number of trials) because it would miss the opportunity to exploit high-value regions. Thus, to obtain a positive control, we simulated an infinitely exploratory agent, a model that selected a random action (time step) on each trial with equal probability. The KF U → V model, which used uncertainty-driven exploration to choose actions early in learning, tended to outperform other models in the first few trials. In addition, models that incorporated uncertainty into choice tended to recover a better estimate of the contingency in the first 10–15 trials than those that chose based on value alone. Finally, whereas models that shifted toward value exploitation later in learning did not improve their approximation of the value function, the pure exploration null agent further refined its estimate.

*Supplementary Figure 4.* The effect of entropy on trial-wise absolute response time changes (RT swings), derived from a multilevel model of all subjects. Predictors of RT swings in the model were: 1) entropy of the value representation, 2) trial, 3) distance of the previously chosen action from the maximum estimated value, 4) value of the chosen action compared to the estimated global maximum value, 5) the magnitude of the previous RT swing, 6) whether the previous outcome was a reward or omission, and 7) distance of the prior RT from the edge of the time interval. Interactions among these predictors were also included in the model, and subject and run were included as random effects. The data depicted represent the model-predicted marginal effect of entropy on RT swings at the mean of all other predictors. Vertical bars adjoining the dots denote the standard error of the predicted value.

*Supplementary Figure 5.* Simulated performance of Frank TC model at different parameter values across contingencies. Response time data were simulated for 500 simulated participants (replications) who completed 500 trials in each of four contingencies: increasing expected value (IEV), decreasing expected value (DEV), constant expected value (CEV), and constant expected value–reversed (CEVR). The order of contingency blocks was randomly permuted across subjects. Response time data were smoothed using a LOESS smoother (span = 10) and averaged across subjects to emphasize general patterns in response times. Parameters in the baseline configuration (top left) were identical to Frank 2009 (Supplementary Material; [2]): *K* = 1500, λ = 0.2,.*v* 0.2, *α*_*G*_ = 0.3, *α*_*N*_ = 0.3, *ρ* = 1000, ε = 3000. As in Frank 2009, -1000ms – 1000ms of random uniform noise was added to each response time. Panel a displays simulations for 50- trial runs; panel b contains simulations for 500 trial runs. In the top right subpanels, *ρ* = 10000, but other parameters are unchanged. In the lower left subpanels, *α*_*N*_ = 1.0, but other parameters are at baseline. In the lower right subpanels, *λ* = 0.5, but other parameters are at baseline.

